# Halogenated cholesterol alters the phase behavior of ternary lipid membranes

**DOI:** 10.1101/2024.09.18.613775

**Authors:** Deeksha Mehta, Elizabeth K. Crumley, Jinchao Lou, Michael D. Best, M. Neal Waxham, Frederick A. Heberle

## Abstract

Eukaryotic plasma membranes exhibit nanoscale lateral lipid heterogeneity, a feature that is thought to be central to their function. Studying these heterogeneities is challenging since few biophysical methods are capable of detecting domains at sub-micron length scales. We recently showed that cryogenic electron microscopy (cryo-EM) can directly image nanoscale liquid-liquid phase separation in extruded liposomes due to its ability to resolve the intrinsic thickness and electron density differences of ordered and disordered phases. However, the intensity contrast between these phases is poor compared to conventional fluorescence microscopy and is thus both a limiting factor and a focal point for optimization. Because the fundamental source of intensity contrast is the spatial variation in electron density within the bilayer, lipid modifications aimed at selectively increasing the electron density of one phase might enhance the ability to resolve coexisting phases. To this end, we investigated model membrane mixtures of DPPC/DOPC/cholesterol in which one hydrogen of cholesterol’s C19 methyl group was replaced by an electron-rich halogen atom (either bromine or iodine). We characterized the phase behavior as a function of composition and temperature using fluorescence microscopy, FRET, and cryo-EM. Our data suggest that halogenated cholesterol variants distribute approximately evenly between liquid-ordered and liquid-disordered phases and are thus ineffective at enhancing the intensity difference between them. Furthermore, replacing more than half of the native cholesterol with halogenated cholesterol variants dramatically reduces the size of membrane domains. Our results reinforce how small changes in sterol structure can have a large impact on the lateral organization of membrane lipids.

## 1. Introduction

It is now widely accepted that lateral lipid heterogeneity is a key feature of eukaryotic plasma membranes (PMs)^1^ even as questions about the nature of these heterogeneities persist.^2^ Given the hundreds of lipid components in the PM, clustering or phase separation is an expected outcome as most pairs of unlike lipids will not have perfectly ideal interactions.^3^ Spatial variation in membrane lipid composition—most notably the preferential association of sphingolipids and cholesterol within clusters termed lipid rafts—can in turn organize membrane-resident proteins based on preferential affinity for raft or non-raft domains.^4^ Despite an abundance of indirect biochemical and spectroscopic evidence that supports the existence of rafts, direct visualization of raft domains in cells has proved elusive owing to their small size and transient lifetimes.^2^ Moreover, teasing apart the influence of specific lipids on domain formation is a daunting task in the complex chemical environment of a cell membrane. Model membrane studies of simplified mixtures have therefore been instrumental in establishing the physicochemical basis for raft formation.^1, 5^

A typical PM raft model combines cholesterol, a low-melting (low-T_M_) unsaturated phospholipid (e.g., DOPC or POPC), and a high-T_M_ lipid (typically fully saturated DPPC, DSPC, or sphingomyelin).^6^ Immiscibility in these mixtures is driven by unfavorable interactions between the saturated and unsaturated phospholipids, which in the absence of cholesterol will separate into coexisting liquid-disordered (Ld) and gel (Lβ) phases. Cholesterol preferentially associates with the high-T_M_ lipid and, at sufficiently high concentration, abolishes long-range order in the gel phase, thus converting it to a liquid phase that nevertheless retains high chain conformational order.^7^ This unique state of matter—the liquid-ordered (Lo) phase—is thought to have properties similar to the raft phase in cell membranes, with the less tightly packed Ld phase serving as a model for the more disordered “sea” that surrounds the raft.^8^

Model membrane studies have provided unique insight into the nature of Ld and Lo phases. Each exhibits the fast lateral lipid diffusion characteristic of fluids,^9^ but Lo phases, enriched in high-T_M_ lipids and cholesterol, have a much higher degree of chain conformational order^10-12^ and are thicker,^13-15^ stiffer,^16^ and less compressible^17^ than Ld phases. While these differential properties are robust and relatively insensitive to precise structural details of the low-T_M_ and high-T_M_ lipids, a remarkable exception is the characteristic domain size when Ld and Lo coexist, which can change by orders of magnitude depending on the identity of the low-T_M_ lipid.^13, 18, 19^ For example, while DOPC invariably produces micron-sized domains that are easily visualized with fluorescence microscopy, POPC and SOPC—with unsaturation in only one acyl chain—do not, though liquid phase separation is still detected by techniques with sub-optical resolution (e.g., FRET, ESR, SANS).^13, 20, 21^ Intriguingly, the vast majority of naturally occurring low-T_M_ lipids possess a fully saturated *sn*-1 chain and one or a few unsaturations in their *sn*-2 chains,^22^ suggesting that such mixed-chain lipids play an important role in regulating raft size in the PM. Still, a mechanistic explanation for these effects remains contentious,^23^ and new experimental tools are needed to distinguish between proposed theoretical models.

Cryogenic electron microscopy (cryo-EM) is among the few techniques capable of imaging nanoscopic phases.^24^ Lipid membranes appear clearly in cryo-EM images as a set of two dark parallel lines,^25^ and we recently showed that the spacing of these lines—which arise from electron density differences between the lipid headgroups and chains—correlates strongly with bilayer thickness.^26^ Moreover, because ordered and disordered membrane phases are distinguished in part by different thickness, phase separation can be directly visualized with cryo-EM. Notably, contrast in cryo-EM images is much weaker than that provided by strongly partitioning fluorescent probes in an optical microscopy experiment.^26, 27^ In principle, cryo-EM contrast could be enhanced with lipids modified to have greater electron density, presuming that the modifications do not perturb the packing density of the surrounding lipids. Brominated phospholipids and sterols are promising candidates and have been successfully used in X-ray scattering^28, 29^ and cryo-EM^30^ experiments to obtain information about the spatial localization of these molecules within the membrane. Because tightly packed ordered phases already have an inherently greater electron density than disordered phases, molecules that localize to the Lo phase (e.g., sterols) are natural targets for selective halogenation with the goal of increasing Lo/Ld contrast.

Considering that a substantial fraction of target molecules would need to be labeled to achieve this goal, it is crucial to assess the influence of lipid halogenation on membrane phase behavior. This is especially important when the target molecule is cholesterol, due both to its great abundance in animal cell plasma membranes and its potency in modulating membrane physical properties.^31^ Seemingly minor changes to sterol structure can significantly influence membrane properties including permeability,^32, 33^ thickness,^34^ area compressibility,^35, 36^ bending rigidity,^36, 37^ and order.^33, 36^ Indeed, few natural or synthetic sterol variants are as effective as cholesterol at ordering saturated lipid chains in the fluid phase and thus inducing Lo phases.^38-41^ Structural perturbations that can alter the ability to form ordered domains include changes to the hydroxyl headgroup,^38, 40, 42^ the addition of polar or bulky moieties to the smooth alpha face of the steroid ring system,^35, 40^ variation in the number or position of double bonds in the steroid rings,^35, 40^ and alterations to the length, degree of saturation, and/or branching of the aliphatic side chain.^35, 39, 43, 44^ These observations underscore that any proposed perturbation to cholesterol’s structure should be vetted for its effect on membrane lateral organization.

Here, we investigated the effect of halogenation at the C19 position of cholesterol. This single-atom modification to the rough beta face of the steroid ring system would appear to be a reasonable candidate for preserving the Ld + Lo phase coexistence region in ternary lipid mixtures and enhancing the contrast between sterol-rich and sterol-poor domains. Contrary to these expectations, we found that replacing the native cholesterol with either brominated or iodinated variants dramatically reduces the size of membrane domains with little or no change in the contrast between ordered and disordered phases.

## 2. Materials and Methods

### 2.1 Materials

Phospholipids 1,2-dioleoyl-sn-glycero-3-phosphocholine (DOPC), 1,2-dipalmitoyl-sn-glycero-3-phosphocholine (DPPC), and 1-palmitoyl-2-oleoyl-sn-glycero-3-[phospho-rac-(1-glycerol)] (sodium salt) (POPG) were purchased from Avanti Polar Lipids (Alabaster, AL). Cholesterol was purchased from Nu-Chek Prep (Elysian, MN). The fluorescent dyes 1,2-dioleoyl-3-[16-N-(Lissamine rhodamine B sulfonyl) amino]palmitoyl-sn-glycerol] (LR-PR), 1-stearoyl(2-NBD)-2-oleoyl-sn-glycero-3-phosphocholine (NBD-DSPE), and 1-palmitoyl-2-(dipyrromethene boron difluoride)undecanoyl-sn-glycero-3-phosphocholine (TFPC) were from Avanti Polar Lipids; Di-4-ANEPPDHQ (Di-4), and 1,1’-dioctadecyl-3,3,3’,3’-tetramethylindodicarbocyanine, 4-chlorobenzenesulfonate salt (DiD) were from ThermoFisher Scientific (Waltham, MA); and naphtho[2,3-a]pyrene] (naphthopyrene, Naph) was from TCI America (Portland, OR). All phospholipids, dyes, and cholesterol were dissolved in HPLC-grade chloroform and stored at - 20°C until use. The concentration of lipids and cholesterol was determined gravimetrically assuming two tightly bound waters for phospholipids. The concentration of fluorescent dyes was determined from absorbance measurements using the following extinction coefficients: 95,000 M^-1^cm^-1^ (LR-PE), 21,000 M^-1^cm^-1^ (NBD-DSPE), 23,800 M^-1^cm^-1^ (Naph), 96,900 M^-1^cm^-1^ (TFPC), 249,000 M^-1^cm^-1^ (DiD), and 36,000 M^-1^cm^-1^ (Di-8 and Di-4). Ultrapure water was obtained from a Milli-Q IC 7000 purification system (Millipore Sigma, Burlington, MA).

### 2.2 Synthesis of halogenated cholesterol

The synthetic routes for Chol-I19 **1.1** and Chol-Br19 **1.2** are shown in Scheme 1. 19-Hydroxyl cholesterol **1.6** was synthesized from cholesteryl acetate **1.3** according to a previously reported protocol^45^ with slight modifications, including the formation of bromohydrin intermediate **1.4**, radical reaction and the intramolecular ether formation of **1.5** and Zn-AcOH mediated reductive ring opening. Tosylation of the hydroxyl group of **1.6** resulted in **1.7**, which was reacted with NaI or NaBr in isopropanol to afford **1.8** and **1.9**, respectively. The final acetyl deprotection reaction with NaOMe resulted in the desired final targets. It is worth noting that when synthesizing **1.8**, we observed the formation of a rearranged byproduct in which the iodomethyl ended up at the C6 position and the double bonds switched to C5/C10. However, this type of rearrangement reaction was not observed for **1.9** synthesis. Full details of the synthesis including NMR data are given in Supporting Information section S1.

### 2.3 Confocal fluorescence microscopy

Giant unilamellar vesicles (GUVs) were prepared by electroformation as previously described.^46^ Briefly, a mixture of lipids (250 nmol total) and dyes (LRPE and NBD-DSPE at probe/lipid ratios of 1/2000 and 1/1000, respectively) diluted in chloroform was spread onto the ends of two ITO-coated glass microscope slides (Delta Technologies, Loveland, CO) on a 55°C hotplate. Residual chloroform was removed by placing the slides in a heated chamber attached to a vacuum pump, with the chamber maintained at low pressure and 50°C for 2 h. A sandwich was then formed from the two slides using an O-ring spacer filled with 100 mM sucrose solution and placed into an aluminum holder heated to 55°C. GUVs were formed by applying a 2 V peak-to-peak, 10 Hz waveform to the sandwich for 2 h while the holder was maintained at 55°C, followed by cooling to 22°C at a rate of 2.7°C per hour. GUVs were harvested from the chamber and stored in plastic microfuge tubes prior to imaging.

Prior to imaging, 50-250 μL of freshly harvested GUVs in 100 mM sucrose were suspended in 2 mL of 100 mM glucose in a culture tube. After ∼ 30 min, a 4 μL aliquot was taken from the bottom of the tube and sandwiched between a glass slide and cover slip for microscopy observations. Imaging was performed with a Nikon C2+ point scanning system attached to a Nikon Eclipse Ti2-E microscope (Nikon Instruments, Melville, NY) equipped with a Plan Apo Lambda 60×/1.4 NA oil immersion objective. A sample temperature of 22°C was maintained with an objective cooling collar (Bioptechs, Butler, PA). NBD-DSPC and LRPE were excited with 488 nm and 561 nm laser lines, respectively, with quarter wave plates (ThorLabs, Newton, NJ) inserted in the excitation path to correct for polarization artifacts.

### 2.4 Cryo-EM imaging

Aqueous lipid dispersions at 3 mg/mL total lipid concentration were prepared by first mixing appropriate volumes of lipid stocks in chloroform with a glass Hamilton syringe. The solvent was evaporated with an inert gas stream and the sample was kept under vacuum overnight. The dry lipid film was hydrated with ultrapure water at 45°C for ∼ 1 h with occasional vortex mixing, and the resulting multilamellar vesicle (MLV) suspension was subjected to five freeze/thaw cycles between a −80°C freezer and a 45°C water bath. Large unilamellar vesicles (LUVs) were produced by extrusion through a 100 nm polycarbonate filter using a handheld mini-extruder (Avanti Polar Lipids) heated to 55°C, with the suspension passed through the filter 31 times. The size and polydispersity of each LUV preparation was assessed using dynamic light scattering (LiteSizer 100, Anton Paar USA, Vernon Hills, IL) immediately after preparation and immediately before cryo-preservation, which was typically performed 2 days after preparation.

To cryo-preserve LUV samples, a 4 μL aliquot was applied to a Quantifoil 2/2 carbon-coated 200 mesh copper grid (Electron Microscopy Sciences, Hatfield, PA) that was glow-discharged for 30 sec at 20 mA in a Pelco Easi-Glow discharge device (Ted Pella, Inc., Redding, CA). After manual blotting at room temperature (∼ 22°C), the grids were plunged into liquid ethane cooled with liquid N_2_. Cryo-preserved grids were stored in liquid N_2_ until use.

Cryo-EM image collection was performed at ∼ 2 μm under focus on a Titan Krios operated at 300 keV equipped with a Gatan K2 Summit direct electron detector operated in counting mode. Data collection was conducted in a semi-automated fashion using Serial EM software operated in low-dose mode.^47^ Briefly, areas of interest were identified visually and 8×8 montages were collected at low magnification (2400×) at various positions across the grid and then individual areas were marked for automated data collection. Data was collected at 2.7 Å/pixel. Movies of 30 dose-fractionated frames were collected at each target site with the total electron dose being kept to < 20 e^−^/Å^2^. Dose-fractionated movies were drift-corrected with MotionCor2.^48^ Defocus and astigmatism were assessed with CTFFind4.^49^ Finally, a high pass filter was applied along with phase flipping using the ‘mtffilter’ and ‘ctfphaseflip’ routines in the IMOD v4.11 software package.

Projection images of vesicles were analyzed in Wolfram Mathematica v. 13 (Wolfram Research Inc., Champaign, IL) as previously described^26^ to obtain spatially resolved intensity profiles (IPs) in the direction normal to the bilayer. Briefly, vesicle contours (i.e., the set of points corresponding to the midplane of the projected bilayer as defined by a relatively bright central peak) were first generated using a neural network-based algorithm (MEMNET) that is part of the TARDIS software package.^50^ The MEMNET contour was resampled at arc length intervals of ∼ 5 nm, resulting in a polygonal representation of the 2D contour. For each polygon face, all pixels within a 5×20 nm rectangular region of interest centered at the face were selected, and their intensities binned at 1 Å intervals in the long dimension (i.e., normal to the face) and subsequently averaged in the short dimension to produce a local segment IP.

To determine the phase state of each bilayer segment, we subjected the local IP vectors to 2-means clustering using Mathematica’s FindClusters algorithm with the distance function set to Euclidean and all other parameters set to their default values. We also calculated the local bilayer thickness, *D*_*TT*_, as the distance between the two minima of the local IP. Two methods were used to locate the minima: (1) a ‘model-free’ method, in which a local 5-point Gaussian smoothing was first performed, and the distance between the two absolute minimum intensity values on either side of the central peak was determined; (2) a ‘model-fit’ method that fits the profile as a sum of four Gaussians and a quadratic background, with the troughs corresponding to the two absolute minimum intensity values on either side of the central peak. The two methods typically agree to within 1 Å, and the *D*_*TT*_ values we report are the average of the two measurements.

### 2.5 Fluorescence spectroscopy

Vesicles of low average lamellarity (LLVs) were prepared using the method of rapid solvent exchange (RSE) as previously described.^51^ First, desired lipid and probe mixtures were prepared by dispensing stock solutions into 13×100 mm glass screw-cap test tubes containing 25 μL of chloroform. 400 μL of ultrapure water (for cryo-EM imaging) or buffer (for FRET and membrane order measurements) was then added to the test tube, which was immediately mounted on the RSE device where it was exposed to vacuum while undergoing vigorous vortexing. During RSE, a slow leak of Ar was introduced into the tube to facilitate removal of chloroform. After 1.5 min under vacuum, the sample was vented to Ar and transferred to a plastic centrifuge tube.

FRET from LLV samples prepared by RSE was measured using a Horiba FluoroLog 3 spectrophotometer (Horiba USA, Irvine, CA). Samples contained three probes comprising two donor/acceptor FRET pairs, namely Naph/DiD and TFPC/DiD. Probe/lipid ratios were 1/200 for Naph, 1/1500 for TFPC, and 1/1000 for DiD. For fluorescence measurements, a 0.100 mL aliquot of a 0.625 mM (total lipid) sample was added to 1.90 mL water in a fluorescence cuvette while stirring at 1000 RPM with a flea stir bar. Fluorescence intensity was measured in five channels (excitation/emission λ in nm): Naph direct signal (427/545); Naph/TFPC FRET (427/664); TFPC direct signal (499/551); TFPC/DiD FRET (499/664); and DiD direct signal (646/664). Excitation and emission slit widths were 3 nm and the signal integration time was 0.1 s. To establish the temperature dependence of FRET, data were collected from 60°C to 10°C in -1°C steps, with the cuvette temperature maintained by a Peltier device (Quantum Northwest, Liberty Lake, WA). FRET was calculated from the intensity measured in the FRET channel after correcting for direct fluorescence contributions from donor and acceptor, using samples that contained only donor or acceptor as controls as described previously.^20^ The miscibility transition temperature, *T*_*misc*_, was obtained by fitting the ratio of the FRET signals from the two probe pairs (i.e., TFPC/DiD FRET divided by Naph/TFPC FRET) to a phenomenological model as described in Supporting Information section S2.

For membrane order measurements, LLVs in aqueous buffer (25 mM HEPES, 150 mM NaCl, 1 mM EDTA) were prepared with the probe Di-4 included at a 1/1000 probe/lipid ratio. Fluorescence emission at 560 nm and 650 nm (I_560_ and I_650_, respectively) was measured using an excitation wavelength of 488 nm for calculating the generalized polarization (GP)^52^:

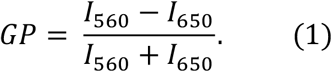

## 3. Results and Discussion

The structures of Chol, Chol-Br19, and Chol-I19 are shown in Fig.1. Our baseline for determining the influence of halogenated cholesterol on cryo-EM intensity and membrane phase behavior was the ternary mixture DPPC/DOPC/Chol, one of the most commonly used and well-studied model systems for lipid rafts. The phase behavior of DPPC/DOPC/Chol has been characterized as a function of composition and temperature by a variety of biophysical techniques including fluorescence microscopy,^6, 11 2^H NMR,^10, 12, 53^ FRET,^54^ wide-angle^55^ and small-angle X-ray scattering,^14^ X-ray diffraction,^56^ and small-angle neutron scattering.^57-59^ For the purposes of this study, the most important features of the room temperature phase diagram are a region of coexisting gel (Lβ) and Ld phases at low cholesterol concentrations (< ∼ 15 mol%), and a region of coexisting Ld and Lo phases at intermediate cholesterol concentrations (∼ 15-35 mol%).

**Figure 1.**
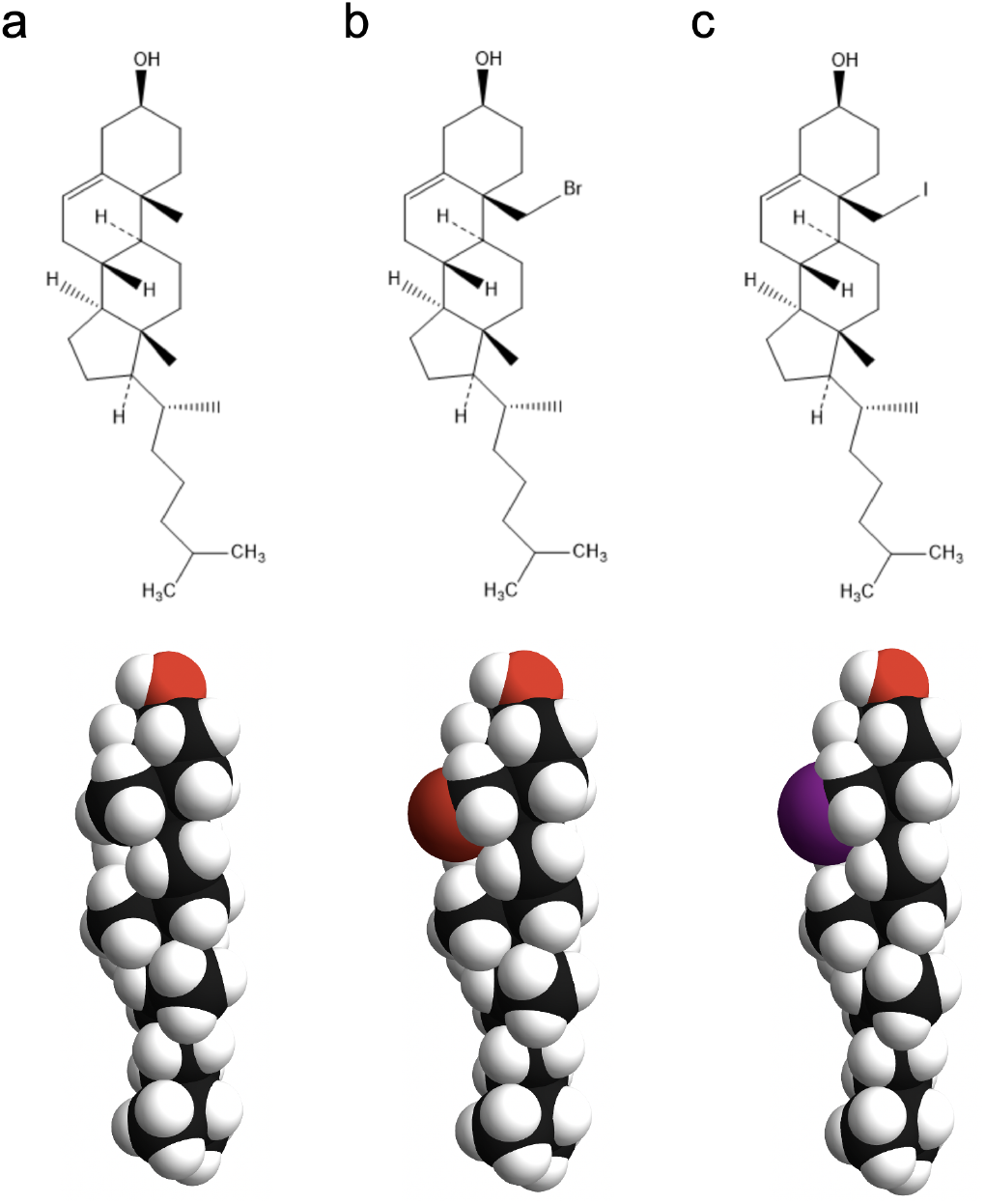
Structures of sterols used in this study. Shown are the chemical structures (top) and space-filling models (bottom) of Chol (a), Chol-Br19 (b), and Chol-I19 (c). Chol-Br19 and Chol-I19 are halogenated analogs of Chol in which one hydrogen at C19 is replaced with a bromine or iodine atom, respectively.

Our results are organized as follows. First, we show how the macroscopic phase behavior visible with fluorescence microscopy changes when Chol is substituted with either Chol-Br19 or Chol-I19. We then determine the nanoscopic mixing behavior of these systems with FRET and cryo-EM. Using unsupervised machine learning to identify Ld and Lo domains in cryo-EM images, we next establish how halogenated cholesterol variants affect the intensity contrast. Finally, we use Di-4 to measure how these variants influence membrane order, leading to a mechanistic explanation for their effects on phase behavior and contrast.

### 3.1 Halogenated sterols eliminate macroscopic Ld + Lo phase separation in GUVs at room temperature

We examined GUVs along compositional trajectories of varying Chol concentration at DPPC:DOPC ratios of 1:2, 1:1, and 2:1 (referred to as trajectories A, B, and C, respectively) and 22°C. Fig. 2 top row shows confocal slices for representative GUVs along trajectory B. Consistent with previous reports,^6^ coexisting Ld + Lβ phases at low Chol concentrations transitioned to Ld + Lo coexistence at higher Chol concentrations, and eventually vesicles of uniform appearance when Chol exceeded 30 mol%. When Chol was replaced by either Chol-Br19 (Fig. 2 middle row) or Chol-I19 (Fig. 2 bottom row), the most striking difference was the absence of macroscopic Ld + Lo phase coexistence at higher sterol concentrations. Although Ld + Lβ phase separation was still visible at low concentrations of halogenated Chol, vesicles with > 10 mol% Chol-Br19 or 15 mol% Chol-I19 were uniform in appearance.

**Figure 2.**
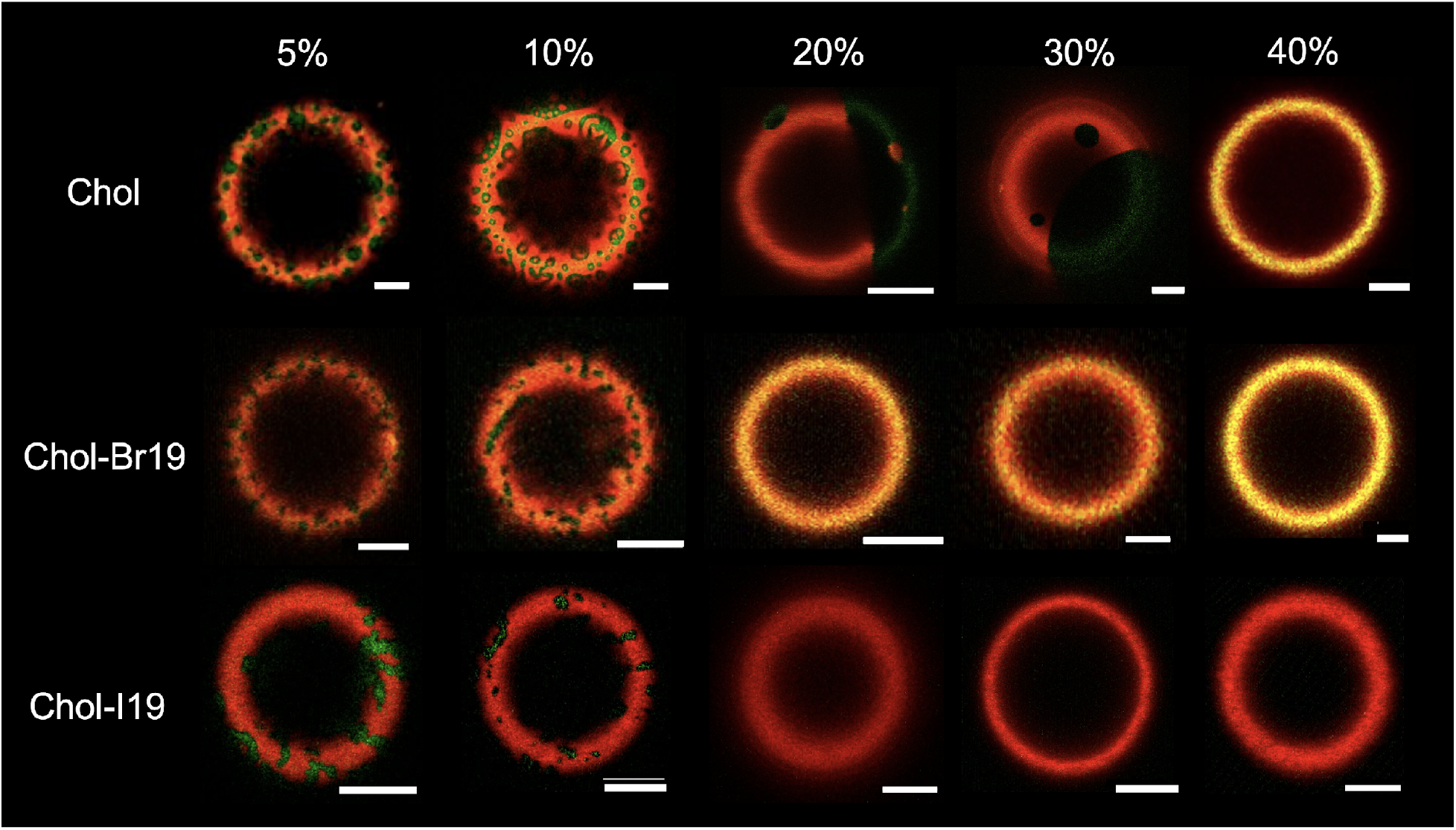
Confocal images of GUVs containing halogenated cholesterol at 22°C. Shown are confocal slices near the vesicle cap for three-component mixtures containing 1:1 DPPC/DOPC plus increasing concentrations of Chol (top row), Chol-Br19 (middle row), or Chol-I19 (bottom row). GUVs were labelled with fluorescent dyes LR-PE (red) and 18:0 NBD (green); the images shown are overlays of the green and red channels. Scale bars are 5 μm.

Fig. 3 summarizes the phase behavior of GUVs for all studied compositions. The ternary phase diagram for DPPC/DOPC/Chol (Fig. 3a) is consistent with literature^6^ and reveals regions of coexisting Ld + Lβ (red squares) and Ld + Lo (green circles). In stark contrast, no Ld + Lo coexistence was observed in either of the halogenated Chol systems at room temperature (Fig. 3b-c). Notably, the phase behavior of GUVs from trajectory A of the Chol-I19 system were difficult to classify at higher Chol-I19 concentrations, with some vesicles showing gel + fluid phases and others appearing to be uniform. We speculate that these inconsistencies may be due to a lower solubility limit for Chol-I19 in DOPC-rich bilayers, but further investigation would be needed to confirm this hypothesis.

**Figure 3.**
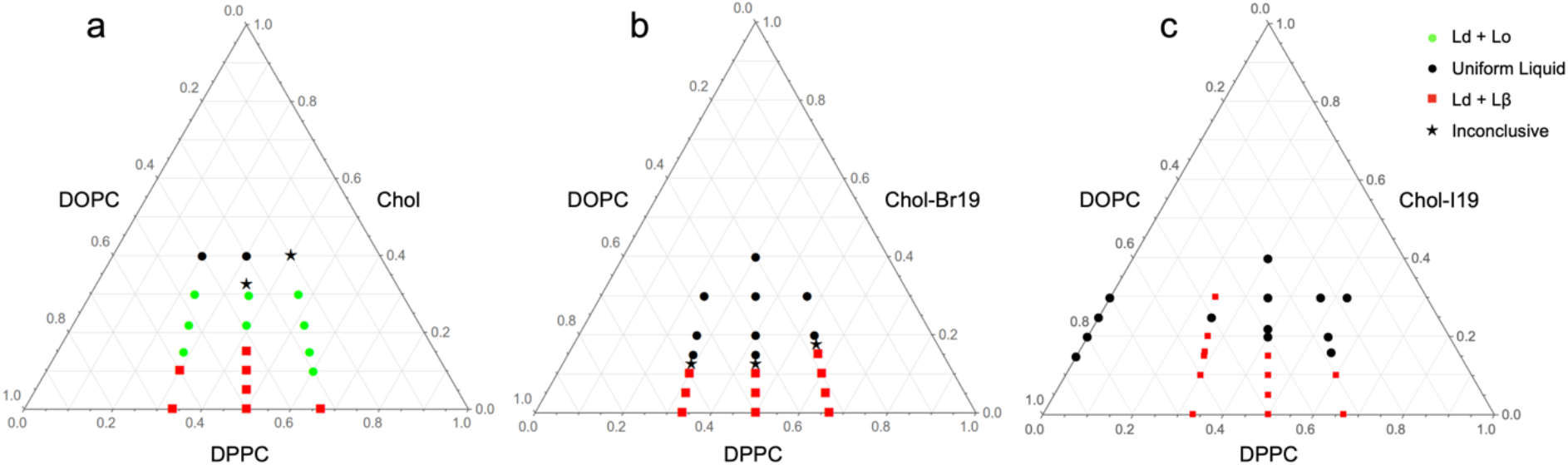
GUV phase diagrams of ternary mixtures containing halogenated cholesterol at 22°C. Ternary phase diagram for a three-component mixture of DPPC/DOPC/Chol (a), DPPC/DOPC/Chol-Br19 (b), and DPPC/DOPC/Chol-I19 (c). Each point shows the predominant phase behavior determined from confocal fluorescence microscopy of GUVs: Ld + Lβ (red squares), Ld + Lo (green circles), or uniform mixing (black circles). Black stars indicate compositions that show approximately equal populations of uniform and phase-separated vesicles (for Chol-containing mixtures) or where small gel domains can no longer be resolved with confidence (for halogenated Chol-containing mixtures).

We next sought to determine the amount of native Chol that could be replaced by Chol-Br19 without disrupting macroscopic Ld + Lo phase coexistence. Fig. 4 reveals that, for the composition DPPC/DOPC/sterol 40/40/20 mol%, Ld + Lo phase separation remained visible in GUVs when as much as one-third of Chol was replaced by Chol-Br19 (Fig. 4a-c); however, replacing two-thirds or more resulted in uniform GUVs (Fig. 4d-e).

**Figure 4.**
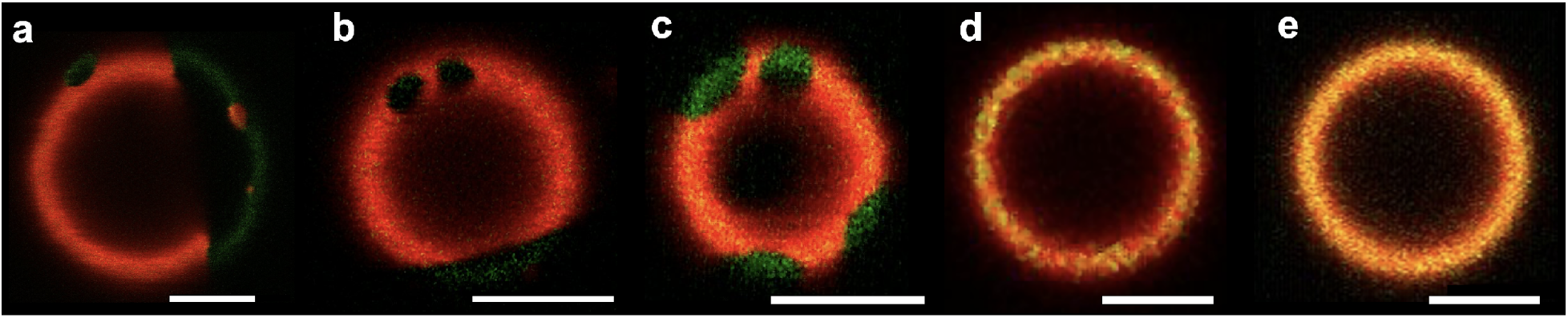
Replacing Chol with Chol-Br19 abolishes macroscopic Ld+Lo phase separation in GUVs at 22°C. Shown are confocal slices near the vesicle cap for three- and four-component mixtures of DPPC/DOPC/Chol/Chol-Br19: 40/40/20/0 mol% (a); 40/40/12.5/7.5 mol% (b); 40/40/10/10 mol% (c); 40/40/7.5/12.5 mol% (d); 40/40/0/20 mol% (e). GUVs were labeled with fluorescent dyes LR-PE (red) and 18:0 NBD (green); the images shown are overlays of the green and red channels. Scale bars are 5 μm.

### 3.2 FRET reveals nanoscopic heterogeneities with lower miscibility transition temperatures in mixtures containing halogenated sterols

It is firmly established that some cholesterol-containing mixtures possess nanoscopic lateral heterogeneity that can be detected by spectroscopic or scattering techniques, but not by conventional fluorescence microscopy.^5, 23, 60^ An especially well-studied case is that of mixtures containing the mixed-chain low-T_M_ lipid POPC and a high-T_M_ lipid species (e.g., DSPC or SM) at intermediate cholesterol concentrations of ∼ 15-30 mol%.^19, 20, 61-63^ Feigenson has termed such mixtures Type I to indicate the presence of a single macroscopic phase coexistence region (i.e., Ld + Lβ) at low Chol concentration, in contrast to Type II mixtures like DPPC/DOPC/Chol that also show macroscopic Ld + Lo phases at higher Chol concentrations.^64^

To assess mixing behavior of the Type I DPPC/DOPC/halogenated-Chol systems at nanoscopic length scales, we measured FRET between lipid fluorophores that are known to show a strong phase preference in the presence of coexisting liquid phases. Specifically, colocalization of donor TFPC and acceptor DiD within Ld phase domains is expected to result in enhanced FRET efficiency, while FRET between Naph donor and DiD acceptor should decrease due to the preference of Naph for Lo phase^21^ and resulting segregation from DiD. Fig. 5a, which shows the ratio of the FRET signal from the two probe pairs as described in Methods, confirms these expectations for the mixture DPPC/DOPC/Chol 40/40/20 mol% (blue circles), revealing an abrupt change in behavior upon cooling through ∼ 37°C. We fit the FRET data to a phenomenological 5-parameter model characterized by a linear regime at high temperature (where the membrane is in a uniform, Ld phase) that switches to a more steeply varying regime below a miscibility transition temperature *T*_*misc*_ that is one of the adjustable parameters (additional details are provided in Supporting Information section S2). The fitted model is shown as a solid blue line in Fig. 5a. The *T*_*misc*_ value obtained for DPPC/DOPC/Chol 40/40/20 mol% (*T*_*misc*_ = 37.1 ± 0.2 °C, indicated by a dashed blue vertical line in Fig. 5a) is higher than the value of 32.5°C obtained by Veatch and Keller from microscopy observations,^65^ consistent with a regime of nanoscale heterogeneity above the macroscopic miscibility transition temperature.

**Figure 5.**
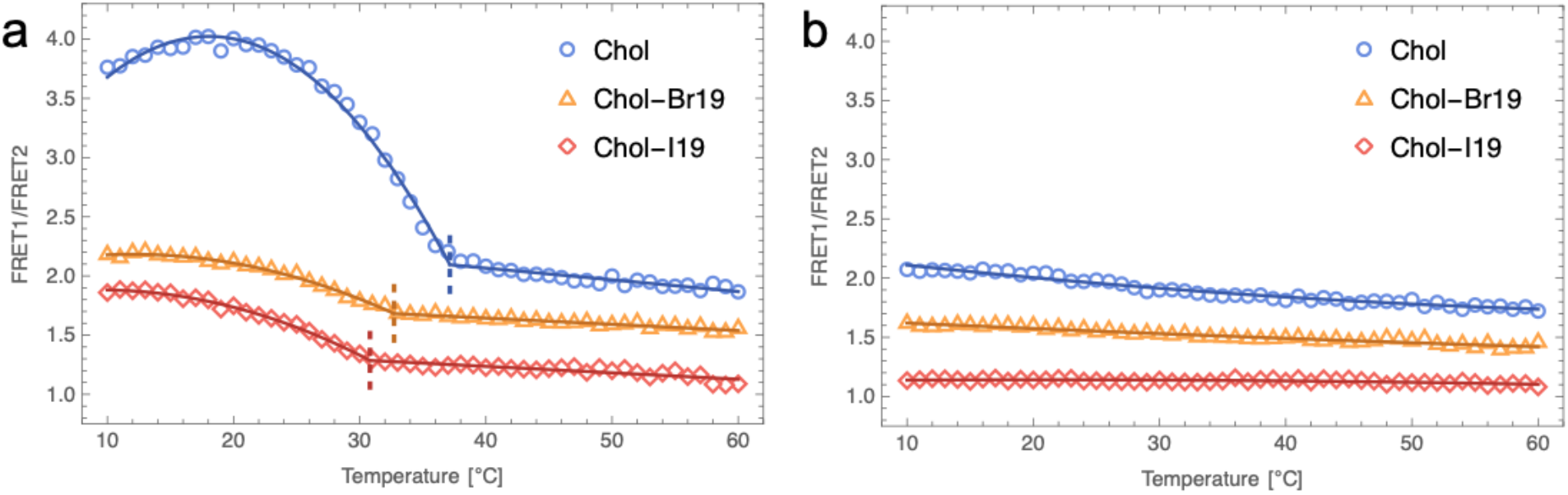
FRET data from ternary mixtures containing halogenated cholesterol. Shown are FRET data vs. temperature for mixtures of 1:1 DPPC/DOPC plus 20 mol% sterol (a) or 40 mol% sterol (b). Solid lines are fits to a model in which the sample has a miscibility transition (for 20 mol% sterol data in panel a) or remains uniformly mixed over the entire temperature range (for 40 mol% sterol data in panel b). The dashed lines in panel a indicate the best-fit value of *T*_*misc*_ for these data. Data from different samples are offset vertically for clarity. Each dataset corresponds to a single sample measured as a function of temperature as described in Methods.

Fig. 5a reveals a similar dependence of FRET on temperature when Chol was replaced with either Chol-Br19 (orange triangles) or Chol-I19 (red diamonds). Although the increase in the FRET ratio below *T*_*misc*_ is attenuated in these systems compared to the Chol-containing system, the data could not be adequately fit by a monotonically varying model for uniform mixing. The solid lines show the best fit to the 5-parameter model and reveal a lowering of *T*_*misc*_ to 32.7 ± 0.3 °C for Chol-Br19 and 30.8 ± 0.3 °C for Chol-I19. The implication of non-random mixing at 22°C in these mixtures is striking given the uniform appearance of GUVs of the same composition and temperature (Fig. 2). For comparison, Fig. 5b shows FRET data for DPPC/DOPC/sterol 30/30/40 and the best fit to a linear model, clearly demonstrating the absence of abrupt changes in FRET in each of these compositions that also have a uniform appearance in GUV images (Fig. 2). *T*_*misc*_ values for all compositions studied are reported in Table S1.

We sought to determine the extent of the nanoscopic miscibility gap in both temperature and composition space with FRET. These data are plotted as contour maps in Fig. 6. For Chol-containing mixtures (Fig. 6 top row), *T*_*misc*_ (indicated by the horizontal portion of the dashed line) increases with increasing DPPC concentration, as does the maximum cholesterol concentration showing heterogeneous mixing (indicated by the vertical portion of the dashed line), in agreement with published phase diagrams.^10^ These heterogeneities clearly persist when Chol is replaced with either Chol-Br19 or Chol-I19 (Fig. 6 middle and bottom rows, respectively), though with markedly smaller changes in FRET that suggest weaker probe partitioning and smaller domains. The region of heterogeneous mixing is also narrower in both composition and temperature: compared to native cholesterol, *T*_*misc*_ is lowered by ∼ 5°C in each trajectory, and compositions with more than 25 mol% halogenated Chol appear to be uniformly mixed at the nanoscale at all temperatures. Importantly, the heterogeneous regions within the dashed lines encompass compositions where GUVs were visibly uniform at 22°C (open circles), suggesting the presence of nanoscopic domains in Chol-Br19 and Chol-I19 mixtures that can be detected by FRET, but not optical microscopy.

**Figure 6.**
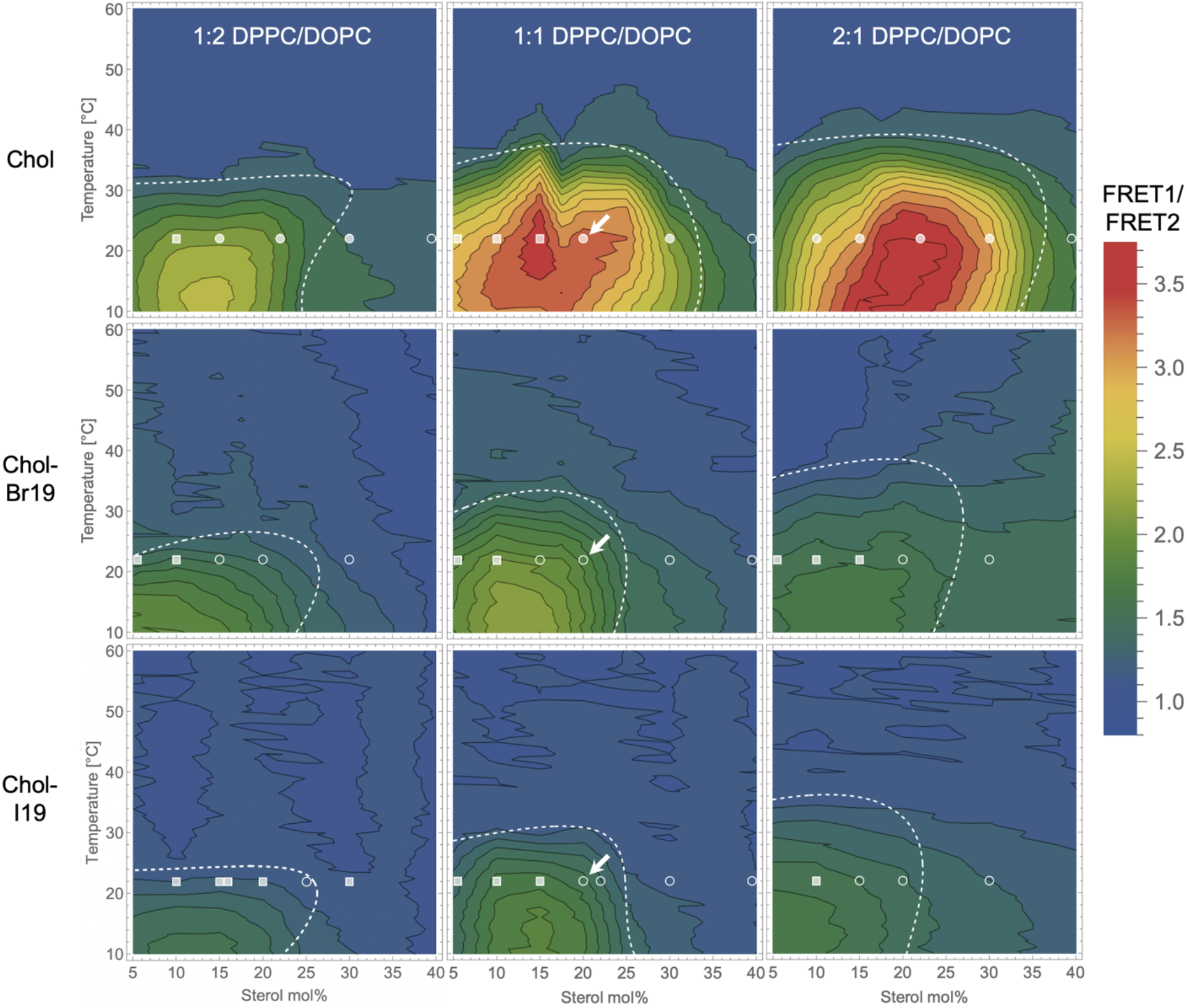
Variation in FRET with temperature and composition. FRET as a function of temperature and sterol mol% is plotted for three DPPC/DOPC ratios (columns) and each of the three sterol variants (rows) as indicated. The dashed lines enclose regions of heterogeneous probe mixing and were determined by fitting horizontal and vertical slices through the surfaces as described in Supporting Information section S2. Symbols denote the macroscopic phase behavior of GUVs imaged at 22°C: Ld + Lβ (filled squares), Ld + Lo (filled circles), uniform (open circles). Arrows denote compositions that were further investigated with cryo-EM.

### 3.3 Halogenated sterols reduce the size of nanodomains in LUVs

We next used cryo-EM to investigate samples where GUVs were uniform, but FRET indicated lateral heterogeneity. We focused on the composition DPPC/DOPC/sterol 40/40/20 mol%, marked with arrows in Fig. 6. For comparison, we also imaged the binary mixtures DOPC/sterol 80/20 mol% that were expected to be uniformly mixed. Representative images of ∼ 100 nm diameter LUVs prepared at these compositions are shown in Fig. 7 and full field-of-view images for the ternary mixtures are shown in Fig. S1.

**Figure 7.**
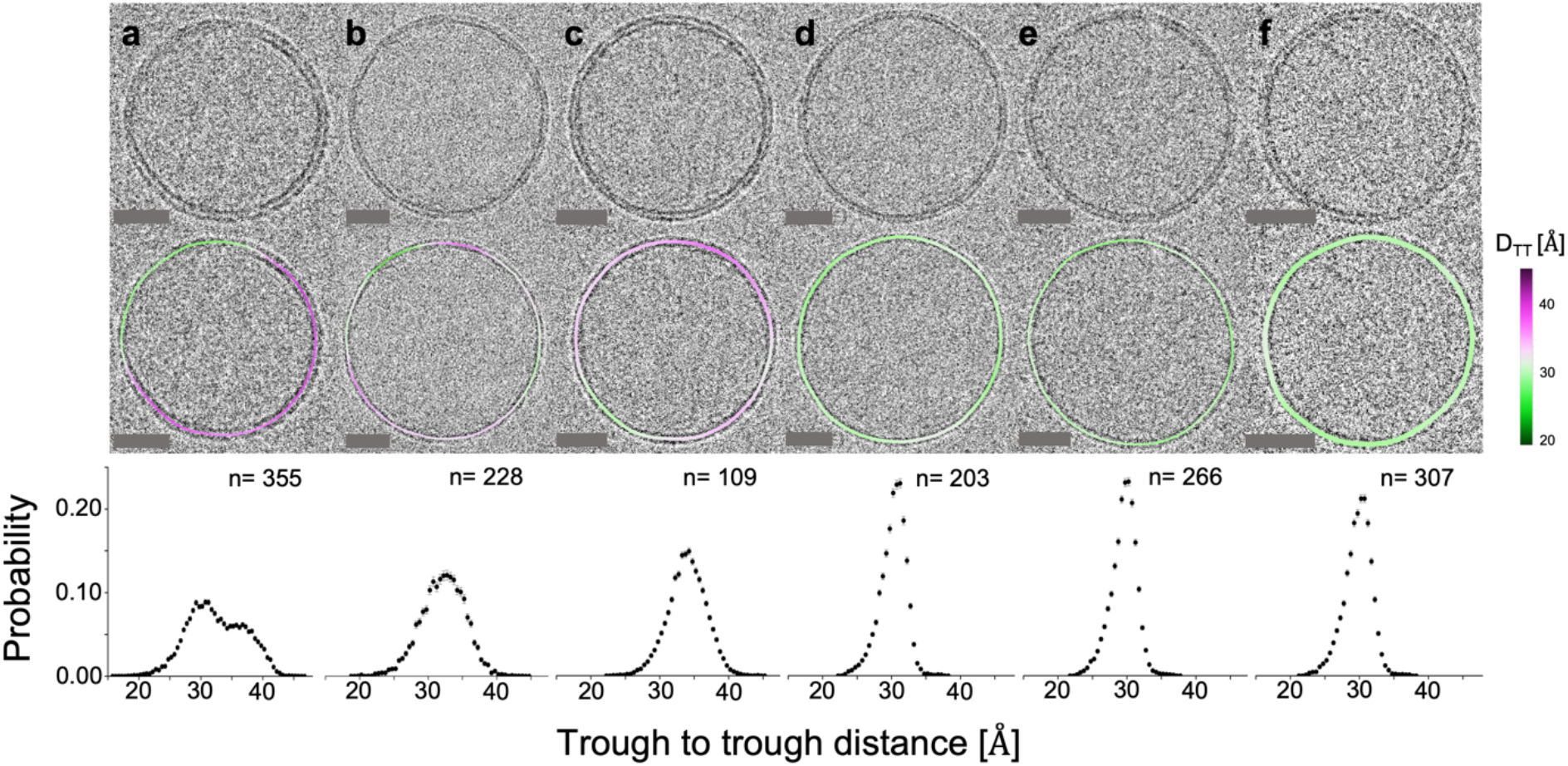
Cryo-EM images of binary and ternary LUVs containing halogenated cholesterol. Vesicle compositions are: DPPC/DOPC/Chol 40/40/20 mol% (a); DPPC/DOPC/Chol-I19 40/40/20 mol% (b); DPPC/DOPC/Chol-Br19 40/40/20 mol% (c); DOPC/Chol 80/20 mol% (d); DOPC/Chol-I19 mol% (e); DOPC/Chol-Br19 mol% (f). Each image is shown without and with a color overlay of the measured bilayer thickness. Scale bars are 25 nm. Plotted in the bottom row are histograms of bilayer thickness distributions obtained from the entire population of vesicles. The number of vesicles analyzed per composition is denoted by *n*.

To assess lateral heterogeneity, the portion of the images corresponding to the bilayer was partitioned into segments of ∼ 5 nm arc length. Intensity profiles (IPs) normal to the bilayer were obtained from each segment and used to determine the local bilayer thickness as previously described.^26^ Consistent with expectations from FRET measurements, each of the ternary mixtures showed substantially greater thickness variation within individual vesicles (Fig. 7a-c) than did any of the binary mixtures (Fig. 7d-f). Most of the DPPC/DOPC/Chol vesicles showed one or a few large domains, while vesicles composed of DPPC/DOPC/Chol-Br19 or DPPC/DOPC/Chol-I19 typically showed several smaller domains. Histograms of segment thicknesses (Fig. 7, bottom row) reveal a clear bimodal distribution for DPPC/DOPC/Chol with peaks centered at ∼ 30 Å and 37 Å corresponding to Ld and Lo phases, respectively. For the ternary mixtures containing Chol-Br19 or Chol-I19, segment thickness distributions were unimodal but substantially broader than distributions obtained from the three binary mixtures. Together, these observations are consistent with lateral thickness heterogeneity in the ternary mixtures that contain halogenated cholesterol, but at a smaller length scale than that found in the ternary mixture with native cholesterol.

To further investigate domain size, we calculated spatial autocorrelation functions (ACFs) of segment thicknesses within individual vesicles as previously described.^25^ The ensemble-averaged ACFs obtained from binary DOPC/sterol mixtures (Fig. 8, upper) were featureless as expected for uniform mixtures, while ACFs from ternary mixtures (Fig. 8, lower) showed clear signatures of lateral heterogeneity. For DPPC/DOPC/Chol, the ACF decayed gradually before reaching a baseline value at a lag distance of ∼ 40 nm, consistent with domain dimensions similar to the vesicle size (i.e., large scale phase separation within the LUV). Remarkably, the ACFs for DPPC/DOPC/Chol-Br19 and DPPC/DOPC/Chol-I19 exhibited a rapid decay and oscillations reminiscent of a nanoemulsion or spatially modulated domain patterning.^66^

**Figure 8.**
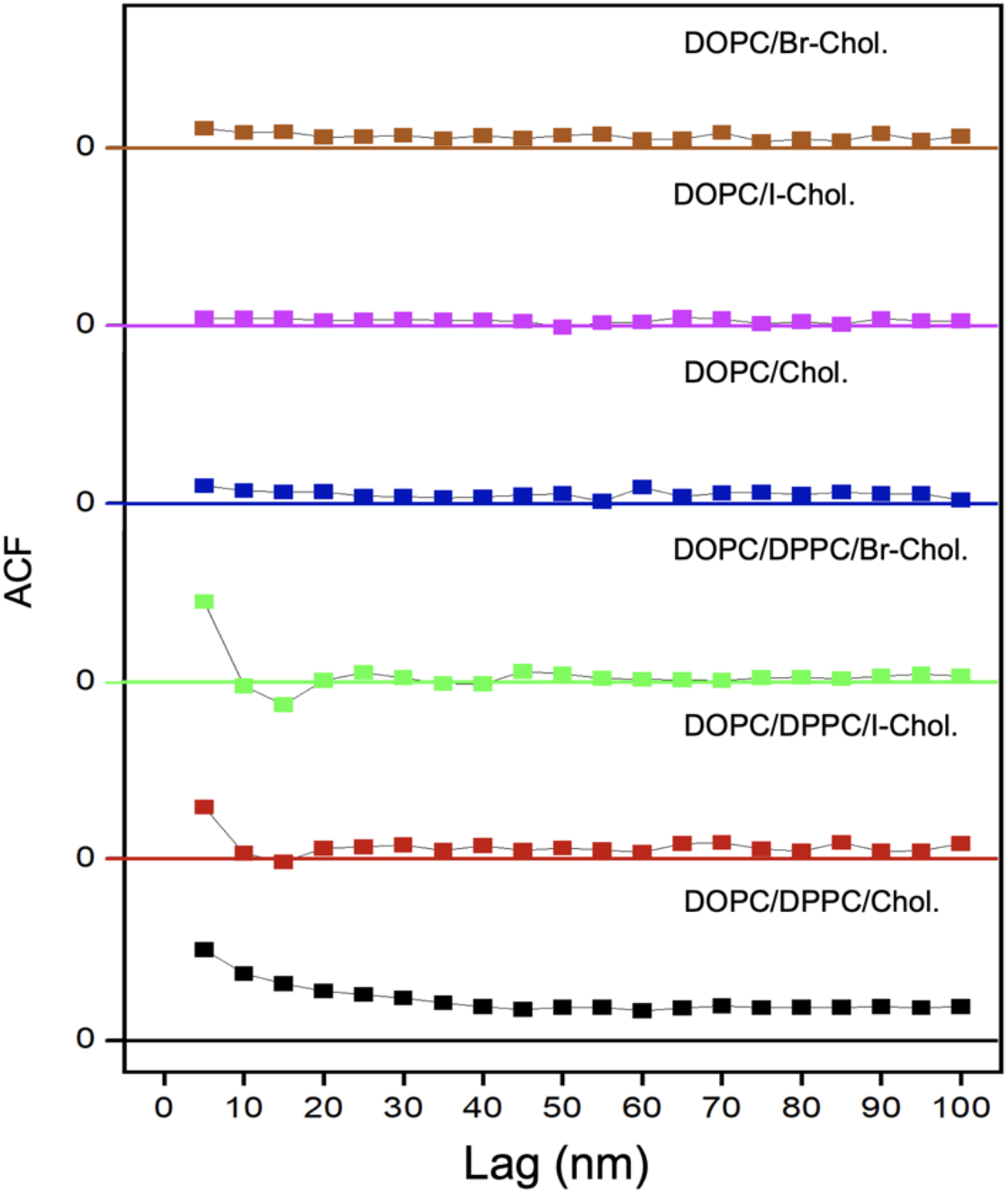
Spatial autocorrelation of bilayer thickness reveals heterogeneity in ternary mixtures. Shown is the ensemble-averaged spatial ACF of bilayer thickness calculated from cryo-EM images for vesicle compositions as indicated. The number of vesicles analyzed for each sample is shown in Table S2.

### 3.4 Halogenated sterols increase the internal intensity contrast of both Ld and Lo phases in cryo-EM images

The primary motivation for this study was to investigate the ability of electron-dense cholesterol analogues to enhance contrast in cryo-EM images of liposomes. To this end, segment IPs obtained from cryo-EM images of DPPC/DOPC/sterol liposomes were subjected to 2-means clustering to group them by their phase state as previously described.^67, 68^ The average IPs of the two clusters, shown in Fig. 9a-b for each of the three sterol variants, show marked differences in both trough-to-trough distances and internal contrasts (which we define as the difference in the maximum and minimum intensity values). Based on these differences, we identify the cluster-1 and cluster-2 segments as belonging to Ld and Lo phases, respectively. The phase fractions and averaged thicknesses obtained from the clustering analysis are provided in Table S2.

**Figure 9.**
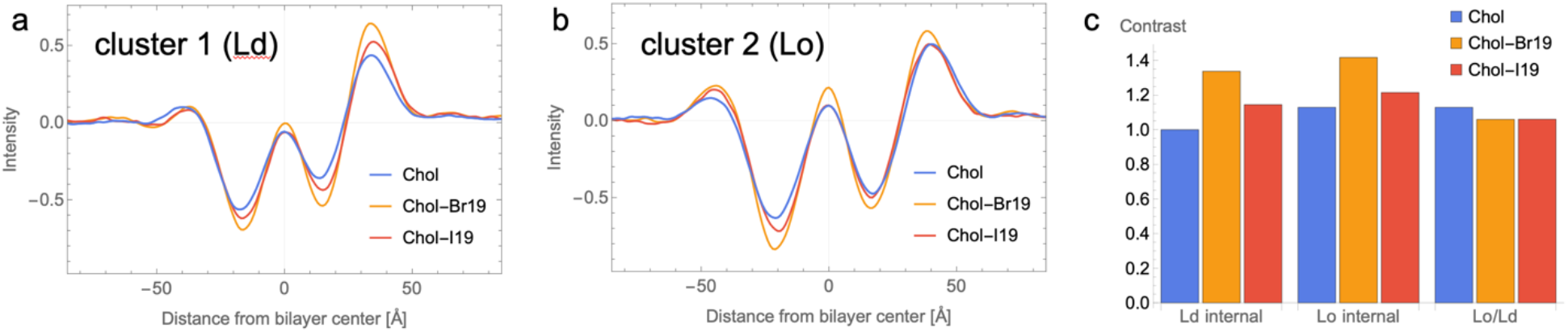
Influence of halogenated cholesterol on cryo-EM intensity contrast. Shown are normalized average intensity profiles (IPs) for the Ld (a) and Lo (b) phases obtained from cryo-EM images of liposomes. Liposome compositions are DPPC/DOPC/Chol 40/40/20 (blue), DPPC/DOPC/Chol-Br19 40/40/20 (orange), or DPPC/DOPC/Chol-I19 40/40/20 (red). Panel c shows the internal contrast of Ld and Lo phases (defined as the intensity difference between the outer peak and the inner trough of the intensity profiles) as well as the Lo/Ld contrast (defined as the internal contrast ratio of the Lo and Ld phases). The number of vesicles analyzed for each sample is shown in Table S2.

Fig. 9c plots the internal contrast of the Ld and Lo phases as well as the ratio of the Lo and Ld contrasts for the three sterol variants. Consistent with our initial hypothesis, the internal contrast of liposomes containing a halogenated cholesterol increased relative to those containing native Chol, likely due to the greater electron density conferred by the halogen atom. Surprisingly, the increase in contrast was similar for Ld and Lo phases, suggesting that halogenated cholesterol molecules distribute approximately equally between Ld and Lo. This behavior is unlike native Chol, which selectively partitions into the Lo phase with Kp ∼ 3.^10^ As a result of the roughly even partitioning, the Lo/Ld contrast ratio slightly *decreased* for each of the halogenated variants relative to native Chol, contrary to initial expectations.

### 3.5 Halogenated sterols are not as efficient as native cholesterol at ordering the membrane

Macroscopic phase separation is correlated with a high line tension and large difference in order between the coexisting phases. Our observations that Chol-Br19 and Chol-I19 lower *T*_*misc*_ and disrupt macroscopic Ld + Lo phase separation suggests that these halogenated sterols are not as effective at ordering saturated chains as native cholesterol. To test this hypothesis, we used the fluorescent probe Di-4 to evaluate membrane packing.^52, 69^ The Di-4 emission spectrum undergoes a progressive red shift in disordered membrane environments, enabling the calculation of an effective order parameter (GP) as the ratio of the fluorescence intensity at two emission wavelengths as described in Methods.

Fig. 10 shows GP values obtained from compositions that are representative of pure Ld and Lo phases. Samples that contained Chol-Br19 or Chol-I19 showed lower GP values (and hence, reduced order) in both phases compared to the sample that contained native Chol. Interestingly, the reduction in order was substantially larger for the Lo phase, consistent with our hypothesis of a smaller order difference between Lo and Ld for samples with halogenated cholesterol variants. It is likely that the bulky halogen atom disrupts the tight packing of saturated chains within the Lo phase, resulting in weaker sterol partitioning. The smaller difference in order likely results in a lower line tension, consistent with our observation of smaller domains for the halogenated sterol mixtures.

**Figure 10.**
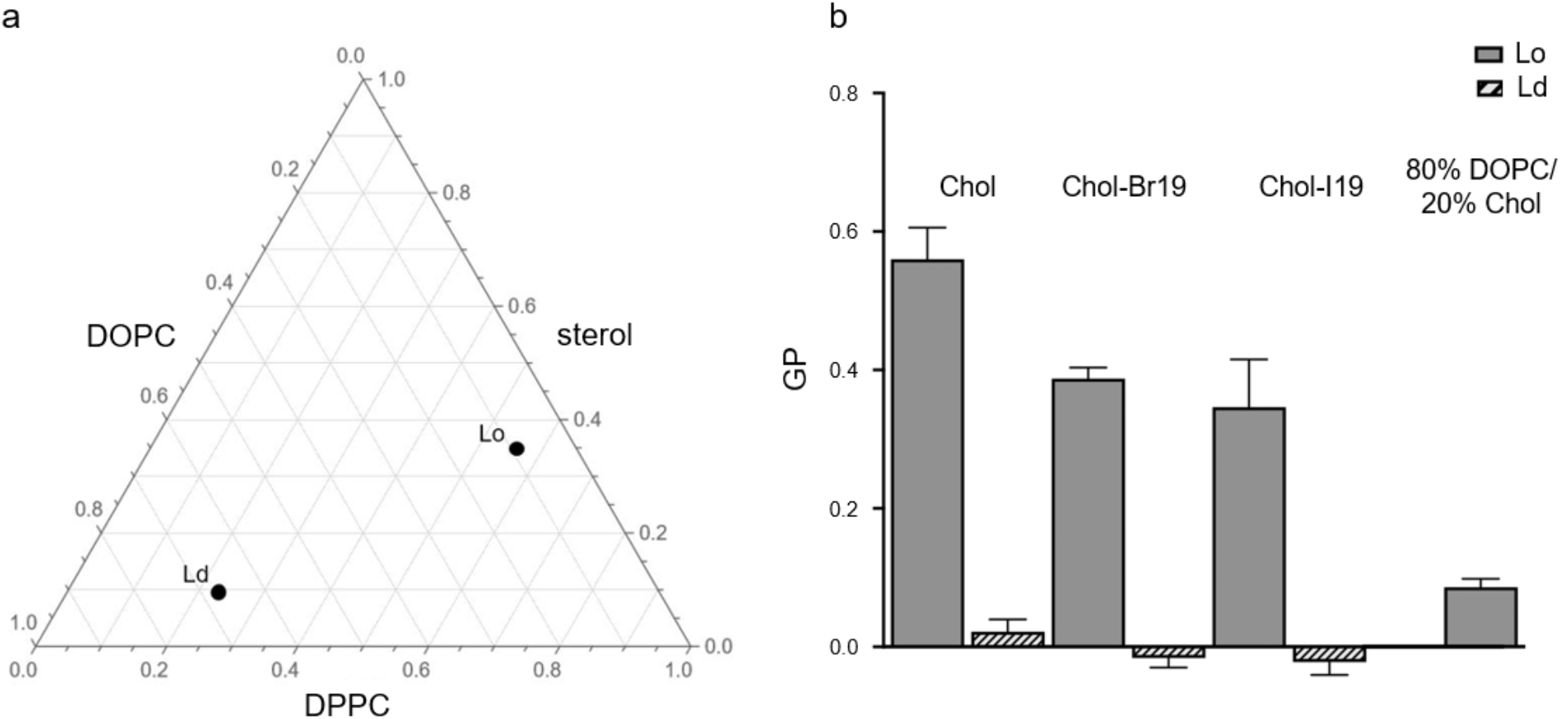
GP values for Lo and Lo phases with different sterols. Representative Ld and Lo compositions are shown in panel a, and GP data for these compositions are shown in panel b for mixtures containing Chol, Chol-Br19, and Chol-I19. GP data for the binary mixture DOPC/Chol 80/20 mol% is also shown for comparison. Error bars are calculated from standard deviation from mean.

## 4. Summary and Conclusion

We investigated the use of halogenated cholesterol analogues to selectively enhance the contrast of cholesterol-rich phases in cryo-EM images of liposomes. We studied ternary mixtures of DPPC/DOPC/Chol, which is frequently used as a model for raft formation in eukaryotic plasma membranes. We found that complete replacement of Chol with either Chol-Br19 or Chol-I19 dramatically alters the macroscopic phase behavior of these systems (Fig. 3) without significantly enhancing the ability to distinguish Lo from Ld in cryo-EM images (Fig. 9). With respect to the phase behavior, replacing native Chol with either of the halogenated variants disrupts macroscopic Ld + Lo domains in GUVs (Fig. 2 and Fig. 4). Instead, the mixtures with halogenated cholesterol form nanoscopic heterogeneities that are detected by FRET (Figs. 5-6) and cryo-EM (Figs. 7-8). Di-4 measurements reveal a large decrease in membrane order for Lo phases, and a smaller decrease for Ld phases, that contain halogenated cholesterol (Fig. 10). This suggests that the bulky halogen atom renders these molecules less efficient at ordering membranes compared to native cholesterol and lessens their preference for the Lo phase. A reduction in membrane order is also consistent with FRET measurements that indicate lower *T*_*misc*_ for mixtures containing halogenated sterols (Table S1). The smaller order difference between Lo and Ld phases offers a potential mechanistic explanation for the formation of nanodomains in mixtures containing halogenated sterols, since the magnitude of the order difference is likely correlated with line tension.^70^

Although Chol-Br19 (and to a lesser extent, Chol-I19) enhanced the internal bilayer contrast at 20 mol% (Fig. 9c), this effect is not specific to the ordered phase because of the roughly even partitioning of halogenated sterols between Lo and Ld. We conclude that the lack of a differential effect on the contrast of coexisting liquid phases, combined with an inability to mimic native cholesterol’s effects on membrane order and phase partitioning, severely limits the usefulness of these molecules for studies of membrane phase behavior. More generally, our results underscore the need to validate cholesterol-like probes for their influence on membrane properties.

## Author contributions

F.A.H. and M.N.W. conceived of the project and obtained funding. J.L. and M.D.B. synthesized reagents. D.M., E.K.C., M.N.W., and F.A.H. designed the research, analyzed the data, and wrote the manuscript. D.M., E.K.C., J.L, M.D.B., M.N.W., and F.A.H. edited the manuscript.

## Acknowledgements

We are grateful to Dr. Tristan Bepler for providing the machine learning algorithm, MEMNET, for automated contouring of vesicles in cryo-EM images, and to Mr. Venkata Mallampali for data acquisition assistance on the Titan Krios. This work was supported by NSF Grant CHE-2204126 (to F.A.H and M.N.W.). M.N.W. acknowledges the William Wheless III Professorship. The Krios was partially supported by CPRIT Core Facility Grant RP190602.

## Figures

**Scheme 1.**
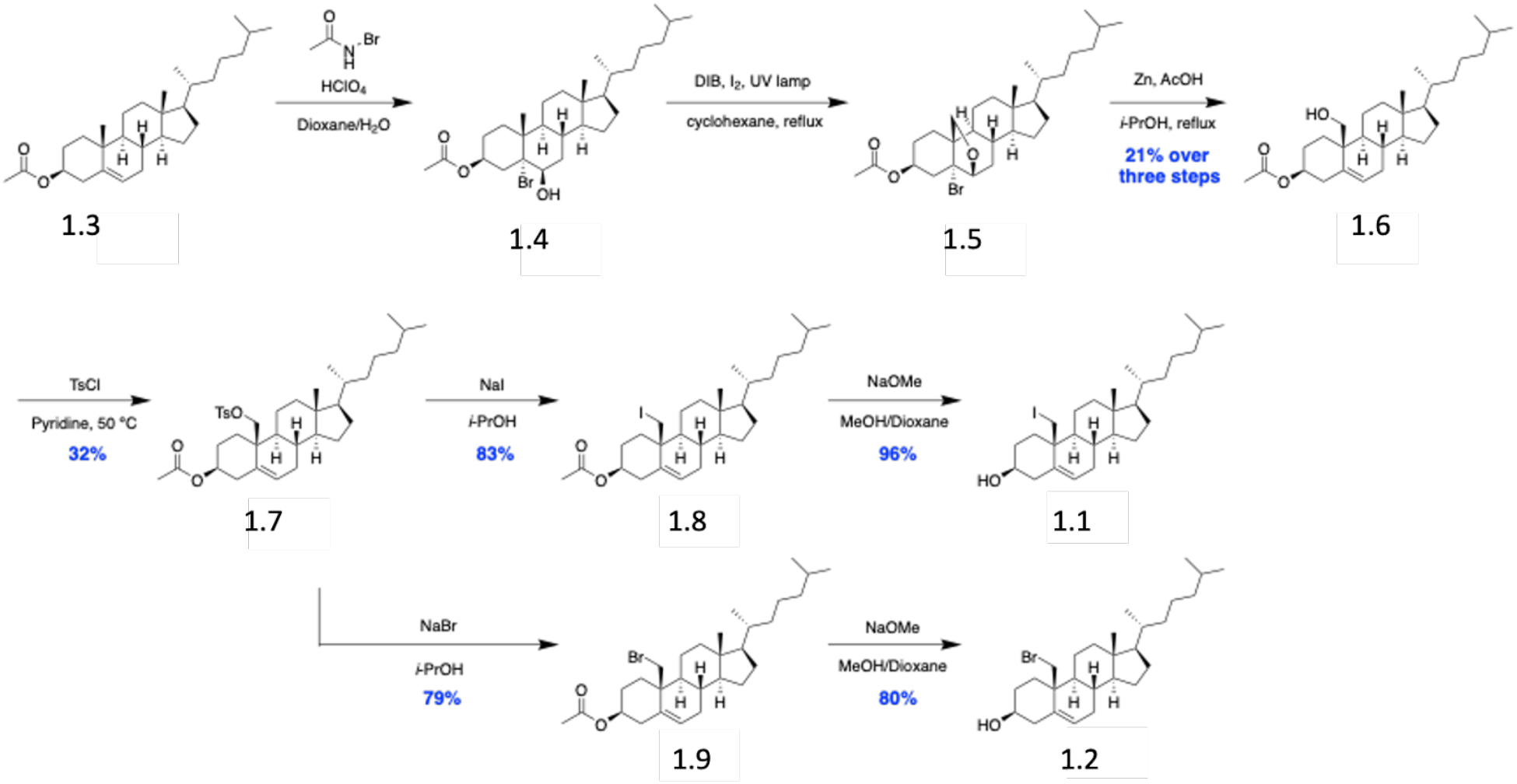
Synthesis of Chol-I19 (1.1) and Chol-Br19 (1.2).

