## Supporting Information for "Halogenated cholesterol alters the phase behavior of ternary lipid membranes"

**This PDF file includes:**

Figure S1
Table S1
Table S2
Sections S1-S2
Supporting References

**Supporting Figures**

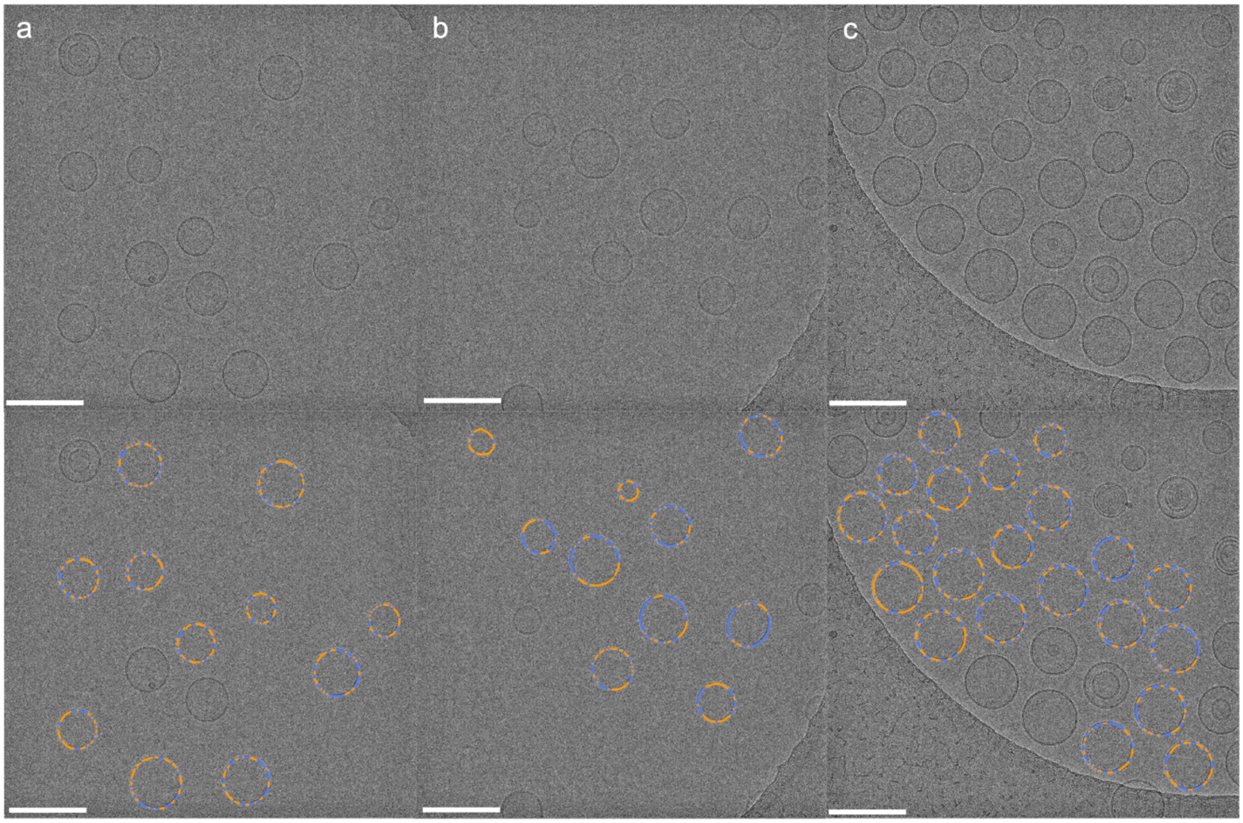


**Figure S1. Full field-of-view cryo-EM images of ternary liposomes.** Vesicle compositions are: DPPC/DOPC/Chol 40/40/20 mol% (a); DPPC/DOPC/Chol-I19 40/40/20 mol% (b); DPPC/DOPC/Chol-Br19 40/40/20 mol% (c). The top row shows the original image, while the bottom row shows the same image with color overlay corresponding to the phase states determined by 2-means clustering as described in the Methods. Scale bars are 200 nm.

### Supporting Tables

**Table S1.** Miscibility transition temperatures ($T_{misc}$) obtained by modeling FRET data from ternary mixtures.

| DPPC:DOPC | Sterol mol% | $T_{misc}$ [°C] Chol | Chol-Br19 | Chol-I19 |
|---|---|---|---|---|
| 1: 2 | 5 | 31.6 | 22.4 | 23.9 |
| | 10 | 31.3 | 27.2 | 25.6 |
| | 15 | 30.6 | 26.7 | 24.4 |
| | 20 | 32.1 | 26.5 | 24.3 |
| | 25 | 33.2 | 23.3 | 24.3 |
| | 30 | 28.0 | 22.6 | 25.2 |
| 1: 1 | 5 | 34.2 | 30.1 | 28.5 |
| | 10 | 36.4 | 33.5 | 32.3 |
| | 15 | 40.4 | 33.2 | 30.5 |
| | 20 | 37.2 | 32.8 | 30.8 |
| | 25 | 36.6 | 31.5 | 31.9 |
| | 30 | 30.7 | 30.1 | 30.5 |
| 2: 1 | 5 | 36.9 | 35.5 | 35.4 |
| | 10 | 38.6 | 37.8 | 36.5 |
| | 15 | 39.5 | 36.1 | 35.5 |
| | 20 | 38.6 | 38.5 | 36.4 |
| | 25 | 36.9 | 35.7 | 35.7 |
| | 30 | 35.3 | 36.2 | 36.2 |

**Table S2.** Number of LUVs analyzed in cryo-EM images.

| *Composition* | | *Number of analyzed vesicles* | *Phase fraction* | | *Phase thickness* (Å) | |
|---|---|---|---|---|---|---|
| Phospholipids | Sterol | | Lo | Ld | Lo | Ld |
| 1:1 DOPC/DPPC | 20% Chol | 355 | 0.45 | 0.55 | 39.03 | 30.79 |
| | 20% Chol-Br19 | 228 | 0.51 | 0.49 | 39.15 | 31.23 |
| | 20% Chol-I19 | 109 | 0.46 | 0.54 | 37.32 | 30.55 |
| 80% DOPC | 20% Chol | 203 | | | 30.2 | |
| | 20% Chol-Br19 | 266 | | | 29.7 | |
| | 20% Chol-I19 | 307 | | | 29.5 | |

#### S1. Synthesis of Chol-Br19 and Chol-I19

The synthetic routes for 19-iodo cholesterol 1.1 and 19-bromo cholesterol 1.2 are shown in Scheme 1. 19-Hydroxyl cholesterol 1.6 was synthesized from cholesteryl acetate 1.3 according to a previously reported protocol with slight modifications,[1] including the formation of bromohydrin intermediate 1.4, radical reaction and the intramolecular ether formation of 1.5 and Zn-AcOH mediated reductive ring opening. Tosylation of the hydroxyl group of 1.6 resulted in 1.7, which was reacted with NaI or NaBr in isopropanol to afford 1.8 and 1.9, respectively. The final acetyl deprotection reaction with NaOMe resulted in the desired final targets. It is worth noting that when synthesizing 1.8, we observed the formation of a rearranged by product, where the iodomethyl ended up at the C6 position and the double bonds switched to C5/C10. However, this type of rearrangement reaction was not observed for 1.9 synthesis.

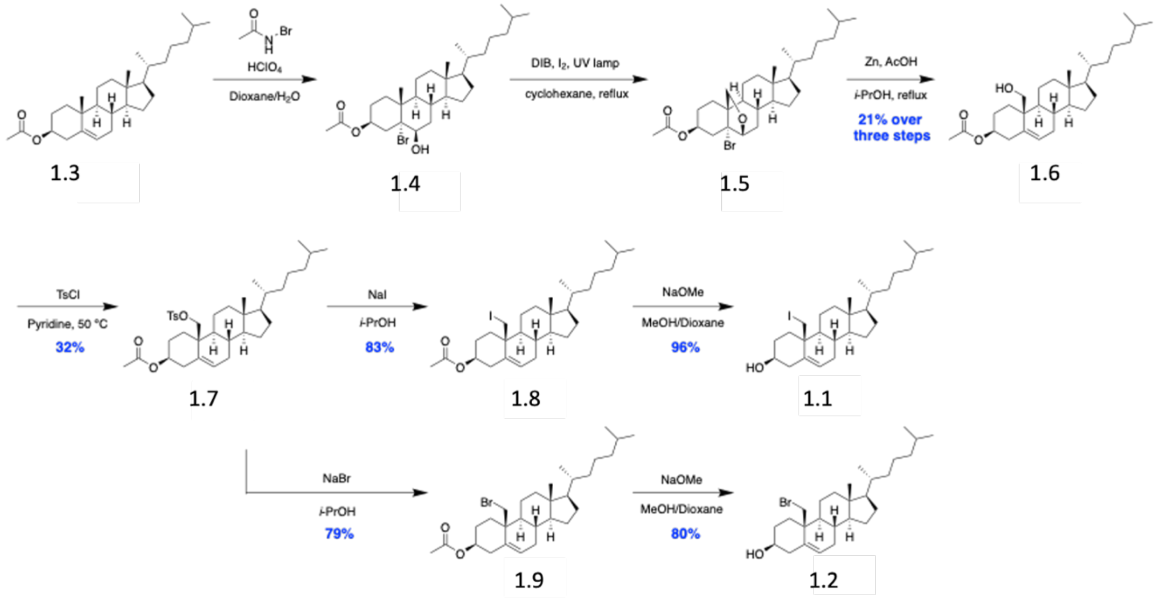


**Scheme1.** Synthetic routes to 19-iodo or 19-bromo cholesterol.

Cholesteryl acetate 1.3 was treated with *N*-bromoacetamide and perchloric acid to form bromohydrin 1.4, which was reacted with iodobenzene diacetate (DIB) and $I_2$ under UV irradiation to form 1.5. Subsequent ring opening followed by tosylation resulted in 1.7, which was treated with NaI or NaBr to introduce the halogens. Final deprotection afforded the 19-iodo cholesterol 1.1 and 19-bromo cholesterol 1.2.

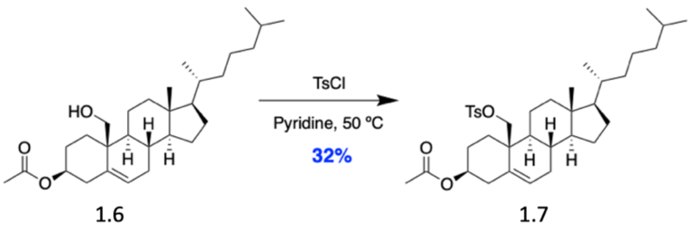


(3*S*,8*S*,9*S*,10*S*,13*R*,14*S*,17*R*)-13-Methyl-17-((*R*)-6-methylheptan-2-yl)-10-((tosyloxy)methyl)-2,3,4,7,8,9,10,11,12,13,14,15,16,17-tetradecahydro-1*H*-cyclopenta[*a*]phenanthren-3-yl acetate (1.7)

Compound 1.6[1] (0.1908 g, 0.429 mmol) was dissolved with 10 mL dry pyridine in a 100 mL RBF under Ar. TsCl (98 mg, 0.514 mmol) was added and the reaction was heated at 50 °C overnight. After removing pyridine under reduced pressure, the crude was redissolved with 50 mL $H_2O$ and 50 mL EtOAc. After collecting the organic layer, the aqueous layer was further extracted with 2 x 25 mL EtOAc. The combined organic layer was further washed with 1 x 100 mL water and 1 x 100 mL brine, respectively, before being dried over $Na_2SO_4$, filtered, and concentrated under reduced pressure. Silica gel column using gradient elution from hexanes to 10% EtOAc in hexanes was needed to afford the product as a white powder (83.6 mg, 0.139 mmol, 32 % yield). $R_f$ = 0.24 (10% EtOAc-hexanes).

$^1$H NMR (500 MHz, $CDCl_3$) δ 7.78 (d, *J* = 8.2 Hz, 2H), 7.34 (d, *J* = 8.1 Hz, 2H), 5.58 (d, *J* = 4.7 Hz, 1H), 4.57 (tt, *J* = 11.3, 4.7 Hz, 1H), 4.10 (d, *J* = 10.2 Hz, 1H), 4.00 (d, *J* = 10.2 Hz, 1H), 2.45 (s, 3H), 2.37 – 2.30 (m, 1H), 2.16 – 2.02 (m, 2H), 2.01 (s, 3H), 1.98 – 1.89 (m, 2H), 1.80 (ddd, *J* = 19.2, 15.2, 9.5 Hz, 2H), 1.56 – 0.94 (m, 21H), 0.93 – 0.81 (m, 12H), 0.53 (s, 3H). $^{13}$C NMR (126 MHz, $CDCl_3$) δ 170.58, 144.87, 133.39, 133.07, 129.94, 128.10, 128.07, 73.09, 69.84, 57.15, 56.15, 49.94, 42.41, 40.11, 39.91, 39.65, 38.09, 36.28, 35.88, 32.54, 32.52, 31.40, 28.26, 28.14, 27.79, 24.16, 23.95, 22.95, 22.69, 21.83, 21.78, 21.47, 18.81, 11.98. HRMS-DART: $[M+NH_4]^+$ calcd for $C_{36}H_{58}O_5SN$: 616.4036; Found: 616.4148.

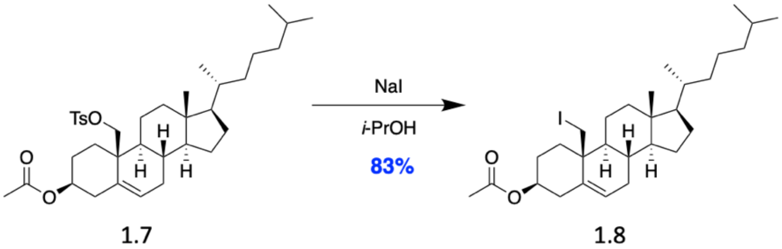


(3*S*,8*S*,9*S*,10*S*,13*R*,14*S*,17*R*)-10-(Iodomethyl)-13-methyl-17-((*R*)-6-methylheptan-2-yl)-2,3,4,7,8,9,10,11,12,13,14,15,16,17-tetradecahydro-1*H*-cyclopenta[*a*]phenanthren-3-yl acetate (1.8)

Compound 1.7 (42 mg, 0.07 mmol) and NaI (22.3 mg, 0.148 mmol) were dissolved with 5 mL isopropanol in a 50 mL RBF. The reaction was heated under reflux without $N_2$ protection for 24 hours. Upon completion, the solvent was removed under reduced pressure and the crude was subjected to silica gel column. Gradient elution from hexanes to 10% EtOAc in hexanes was used to afford the product as a white solid (32 mg, 0.058 mmol, 83% yield). $R_f$ = 0.55 (10% EtOAc-hexanes).

Note: Minor rearranged by product was separated from the column, which has an identical Rf as the product.

$^1$H NMR (500 MHz, $CDCl_3$) δ 5.67 – 5.62 (m, 1H), 4.60 (ddd, *J* = 16.4, 11.4, 4.9 Hz, 1H), 3.57 (d, *J* = 11.0 Hz, 1H), 3.31 (d, *J* = 10.9 Hz, 1H), 2.40 (ddd, *J* = 13.1, 5.0, 2.1 Hz, 1H), 2.15 (ddd, *J* = 14.0, 10.6, 2.5 Hz, 2H), 2.10 – 1.98 (m, 7H), 1.95 – 1.63 (m, 4H), 1.63 – 1.45 (m, 7H), 1.45 – 1.22 (m, 8H), 1.21 – 1.04 (m, 10H), 1.03 – 0.88 (m, 7H), 0.86 (dt, *J* = 6.5, 2.2 Hz, 8H), 0.77 (s, 3H). $^{13}$C NMR (126 MHz, $CDCl_3$) δ 170.61, 135.46, 126.74, 73.21, 57.75, 56.20, 51.27, 42.75, 40.01, 39.67, 38.66, 38.18, 37.30, 36.32, 35.93, 31.96, 31.22, 28.36, 28.16, 28.01, 24.27, 23.97, 22.96, 22.71, 22.23, 21.51, 18.84, 12.44, 11.20.

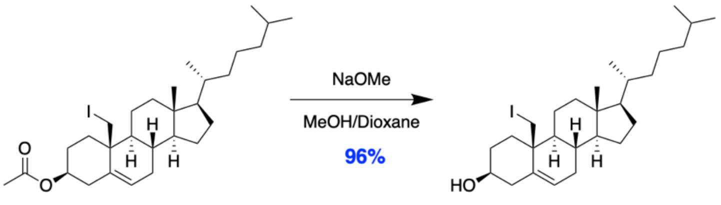


1.8 1.1

(3*S*,8*S*,9*S*,10*S*,13*R*,14*S*,17*R*)-10-(Iodomethyl)-13-methyl-17-((*R*)-6-methylheptan-2-yl)-2,3,4,7,8,9,10,11,12,13,14,15,16,17-tetradecahydro-1*H*-cyclopenta[*a*]phenanthren-3-ol (1.1)

Compound 1.8 (17.5 mg, 0.0316 mmol) was dissolved with 5 mL dioxane/MeOH (1:1, v/v) in a 50 mL RBF at rt. NaOMe solution (4.375 M in MeOH) was next added dropwise until the reaction mixture turned basic, as indicated by pH paper. The reaction progress was monitored through TLC techniques and completed within one hour. After being quenched by addition of 1 N HCl dropwise, the solvent was removed under reduced pressure and the crude was purified via a silica gel column using 25% EtOAc in hexanes. The product was obtained as a white solid (15.5 mg, 0.03 mmol, 96% yield). $R_f$ = 0.44 (25% EtOAc-hexanes).

$^1$H NMR (500 MHz, $CDCl_3$) δ 5.65 – 5.58 (m, 1H), 3.61 – 3.48 (m, 2H), 3.30 (d, *J* = 10.9 Hz, 1H), 2.36 (ddd, *J* = 13.0, 4.9, 2.3 Hz, 1H), 2.08 (dddd, *J* = 21.7, 17.8, 9.0, 5.6 Hz, 5H), 1.90 – 1.65 (m, 4H), 1.59 – 0.97 (m, 24H), 0.95 – 0.82 (m, 12H), 0.77 (s, 3H). $^{13}$C NMR (126 MHz, $CDCl_3$) δ 136.50, 125.75, 71.35, 57.84, 56.23, 51.33, 42.77, 42.34, 40.06, 39.67, 38.59, 37.55, 36.33, 35.93, 31.99, 31.85, 31.22, 28.37, 28.16, 24.28, 23.98, 22.96, 22.71, 22.27, 18.84, 12.45, 11.53. HRMS-DART: $[M+NH_4]^+$ calcd for $C_{27}H_{49}INO$: 530.2859; Found: 530.2890. $[M+H-H_2O]^+$ calcd for $C_{27}H_{44}I$: 495.2488; Found: 495.2498.

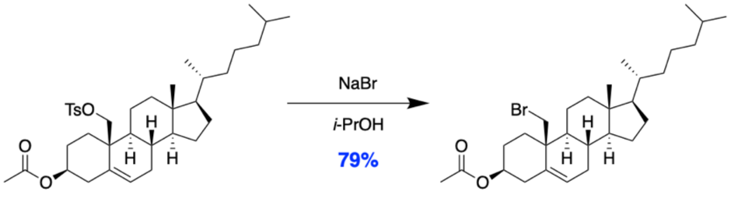


1.7 1.9

(3*S*,8*S*,9*S*,10*S*,13*R*,14*S*,17*R*)-10-(Bromomethyl)-13-methyl-17-((*R*)-6-methylheptan-2-yl)-2,3,4,7,8,9,10,11,12,13,14,15,16,17-tetradecahydro-1*H*-cyclopenta[*a*]phenanthren-3-yl acetate (1.9)

Compound 1.7 (20 mg, 0.0334 mmol) and NaBr (6.87 mg, 0.0668 mmol) were dissolved with 5 mL isopropanol in a 50 mL RBF. The reaction was heated under reflux without $N_2$ protection for 24 hours. Upon completion, the solvent was removed under reduced pressure and the crude was subjected to silica gel column. Gradient elution from hexanes to 10% EtOAc in hexanes was used to afford the product as a white solid (13.4 mg, 0.0264 mmol, 79% yield). $R_f = 0.55$ (10% EtOAc-hexanes). Note: No rearranged by product was observed in this case.

$^1$H NMR (500 MHz, $CDCl_3$) δ 5.67 (dd, *J* = 4.9, 2.5 Hz, 1H), 4.63 (ddd, *J* = 11.5, 6.6, 4.8 Hz, 1H), 3.74 (d, *J* = 11.2 Hz, 1H), 3.51 (d, *J* = 11.2 Hz, 1H), 2.46 – 2.38 (m, 1H), 2.26 – 2.13 (m, 3H), 2.10 – 1.99 (m, 6H), 1.96 – 1.76 (m, 4H), 1.72 (dd, *J* = 13.1, 4.0 Hz, 1H), 1.55 (dddd, *J* = 24.9, 16.9, 8.3, 3.3 Hz, 6H), 1.41 – 1.30 (m, 4H), 1.30 – 1.21 (m, 4H), 1.21 – 0.96 (m, 11H), 0.96 – 0.87 (m, 8H), 0.87 – 0.84 (m, 5H), 0.75 (s, 3H). $^{13}$C NMR (126 MHz, $CDCl_3$) δ 170.62, 134.75, 127.30, 73.27, 57.62, 56.22, 51.01, 42.68, 40.09, 40.04, 39.67, 38.15, 36.32, 36.15, 36.00, 35.93, 32.45, 31.35, 28.36, 28.16, 28.00, 24.27, 23.98, 22.96, 22.71, 22.15, 21.51, 18.84, 12.34. HRMS-DART: $[M-CH_3CO_2]^-$ calcd for $C_{27}H_{44}Br$: 447.2626; Found: 447.2796.

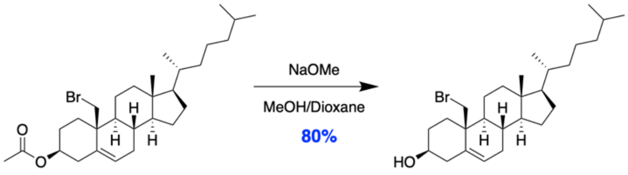


1.9 1.2

(3*S*,8*S*,9*S*,10*S*,13*R*,14*S*,17*R*)-10-(Bromomethyl)-13-methyl-17-((*R*)-6-methylheptan-2-yl)-2,3,4,7,8,9,10,11,12,13,14,15,16,17-tetradecahydro-1*H*-cyclopenta[*a*]phenanthren-3-ol (1.2)

Compound 1.9 (13.4 mg, 0.0264 mmol) was dissolved with 4 mL dioxane/MeOH (1:1, v/v) in a 50 mL RBF at rt. NaOMe solution (4.375 M in MeOH) was next added dropwise until the reaction mixture turned basic, as indicated by pH paper. The reaction progress was monitored through TLC techniques and completed within one hour. After being quenched by addition of 1 N HCl dropwise, the solvent was removed under reduced pressure and the crude was purified via a silica gel column using gradient elution from hexanes to 25% EtOAc in hexanes. The product was obtained as a white solid (9.8 mg, 0.021 mmol, 80% yield). $R_f = 0.4$ (25% EtOAc-hexanes).

$^1$H NMR (500 MHz, $CDCl_3$) δ 5.66 – 5.62 (m, 1H), 3.74 (d, *J* = 11.2 Hz, 1H), 3.56 (ddd, *J* = 16.0, 11.2, 4.9 Hz, 1H), 3.50 (d, *J* = 11.1 Hz, 1H), 2.39 (ddd, *J* = 13.1, 4.9, 2.3 Hz, 1H), 2.14 (tt, *J* = 10.2, 2.7 Hz, 2H), 2.09 – 1.99 (m, 2H), 1.99 – 1.67 (m, 3H), 1.61 – 1.43 (m, 6H), 1.43 – 0.92 (m, 16H), 0.92 – 0.84 (m, 10H), 0.75 (s, 3H). $^{13}$C NMR (126 MHz, $CDCl_3$) δ 135.76, 126.33, 71.41, 57.73, 56.24, 51.09, 42.70, 42.31, 40.10, 40.02, 39.67, 36.38, 36.33, 36.28, 35.93, 32.51, 31.84, 31.35, 28.37, 28.17, 24.28, 23.98, 22.97, 22.71, 22.19, 18.84, 12.35. HRMS-DRAT: [M+H-$H_2O$] calcd for $C_{27}H_{44}Br$: 447.2626; Found: 447.2774. [M+H-HBr] calcd for $C_{27}H_{45}O$: 385.3470; Found: 385.3652.

### S2. Phenomenological model for fitting FRET data

Our FRET metric is sensitized acceptor emission ($F_A^{D_{ex}}$), i.e., we monitor changes in the fluorescence emission of the acceptor under conditions of donor excitation. Because the FRET measurement channel generally also contains contributions from direct donor and acceptor emission, we normalize the FRET signal by the geometric mean of the direct donor ($F_D^{D_{ex}}$) and acceptor ($F_A^{A_{ex}}$) emission, each measured using a combination of excitation and emission wavelengths selective for that probe:

$$FRET = \frac{F_A^{D_{ex}}}{\sqrt{F_D^{D_{ex}} \cdot F_A^{A_{ex}}}}. \qquad (S1)$$

In a uniformly mixed bilayer, the normalized FRET signal typically shows a gradual, monotonic variation with temperature or composition. However, the crossing of a phase boundary will generally be accompanied by an abrupt change in FRET as the fluorescent donor and acceptor lipids either segregate or co-localize in response to lipid demixing.[2] Considering the probes used in this study, TFPC and DiD each partitions favorably into the Ld phase, resulting in increased TFPC→DiD FRET intensity upon Ld + Lo phase separation. Conversely, Nap prefers the Lo phase, leading to reduced Nap→DiD FRET in a phase coexistence regime. When measured as a function of temperature or composition, the FRET signal will change slope upon crossing a phase boundary. In principle, each probe pair should be sensitive to $T_{misc}$. The onset of these changes are, however, often subtle, but can be enhanced by taking the ratio of the FRET signals given by Eq. S1:

$$FRET\ ratio = \frac{FRET_{TFPC\to DiD}}{FRET_{Nap\to DiD}}. \qquad (S2)$$

In practice, we find that the FRET ratio gives a more sensitive indication of $T_{misc}$ than either individual FRET signal. To determine the miscibility transition temperature, $T_{misc}$, we modeled the $FRET\ ratio$ vs. $T$ data with an empirical function of five adjustable parameters that describes two regimes of different behavior:

$$FRET(T; T_{misc}) = \begin{cases} a\left(T^2 - T_{misc}{}^2\right) + b(T - T_{misc}) + FRET_{misc} & T < T_{misc} \\ c(T - T_{misc}) + FRET_{misc} & T \geq T_{misc} \end{cases}. \qquad (S3)$$

In Eq. S3, the parameters $a$ and $b$ control the shape of the curve in the regime of phase separation (i.e., $T < T_{misc}$), the parameter $c$ describes the slope in the uniform regime (i.e., $T \geq T_{misc}$), and the parameter $FRET_{misc}$ is the FRET value at the point where these regimes intersect (i.e., $T = T_{misc}$).
